# FAM104 proteins promote the nuclear localization of p97/VCP

**DOI:** 10.1101/2023.07.25.550451

**Authors:** Maria Körner, Susanne Meyer, Gabriella Marincola, Maximilian Kern, Clemens Grimm, Christina Schülein-Völk, Utz Fischer, Kay Hofmann, Alexander Buchberger

## Abstract

The ATPase p97 (also known as VCP, Cdc48) has crucial functions in a variety of important cellular processes such as protein quality control, organellar homeostasis and DNA damage repair, and its de-regulation is linked to neuro-muscular diseases and cancer. p97 is tightly controlled by numerous regulatory cofactors, but the full range and function of the p97–cofactor network is unknown. Here, we identify the hitherto uncharacterized FAM104 proteins as a conserved family of p97 interactors. FAM104 proteins bind p97 directly *via* a novel, alpha-helical motif and associate with the p97- UFD1-NPL4 complex in cells. FAM104 proteins localize to the nucleus and promote both the nuclear import and chromatin binding of p97. Loss of FAM104 proteins results in slow growth and hypersensitivity to p97 inhibition in the absence and presence of DNA damage, suggesting that FAM104 proteins are critical regulators of nuclear p97 functions.

## Introduction

The abundant, highly conserved AAA+-type ATPase p97 (also known as VCP in mammals and Cdc48 in plants and lower eukaryotes) plays a central role in the maintenance of protein and organelle homeostasis as well as genome integrity (reviewed in Ahlstedt *et al*, 2022, Franz *et al*, 2016a, Papadopoulos & Meyer, 2017). The best-characterized function of p97 is the segregation of ubiquitin-modified proteins in various protein quality control pathways for their subsequent degradation by the 26S proteasome (Brandman *et al*, 2012, Dantuma & Hoppe, 2012, Tanaka *et al*, 2010, Ye *et al*, 2001). p97 is also involved in the lysosomal degradation of endocytic cargo (Ritz *et al*, 2011) and in the selective autophagy of protein aggregates, stress granules as well as damaged mitochondria and lysosomes (Buchan *et al*, 2013, Ju *et al*, 2008, Kim *et al*, 2013, Papadopoulos *et al*, 2017, Tanaka *et al*, 2010, Turakhiya *et al*, 2018). Moreover, p97 possesses a number of crucial nuclear functions, for instance in DNA replication and DNA damage repair (Acs *et al*, 2011, Davis *et al*, 2012, Franz *et al*, 2011, Maric *et al*, 2014, Meerang *et al*, 2011, Moreno *et al*, 2014, Mosbech *et al*, 2012, Mouysset *et al*, 2008, Raman *et al*, 2011). Importantly, mutational perturbation of p97 function causes the neuro-muscular degenerative disease multisystem proteinopathy 1 (MSP1) (Johnson *et al*, 2010, Pfeffer *et al*, 2022, Watts *et al*, 2004), and several cancers and viruses rely on p97 activity, making p97 an attractive target for therapeutic intervention (Das & Dudley, 2021, Deshaies, 2014, Huryn *et al*, 2020).

The molecular basis underlying the diverse cellular functions of p97 is the partial or complete unfolding of substrate proteins by threading through the ring-shaped p97 homohexamer in an ATP hydrolysis-driven process (Blythe *et al*, 2017, Bodnar & Rapoport, 2017, Buchberger, 2022). Since p97 itself lacks appreciable specificity for its physiological substrates, a large number of cofactor proteins control substrate binding, subcellular localization and oligomeric state of p97 (reviewed in Buchberger *et al*, 2015). Some of these cofactors bind to p97 in a mutually exclusive manner and define functionally distinct, major p97 complexes: A heterodimer of UFD1 (also known as UFD1L) and NPL4 (also known as NPLOC4) recruits ubiquitylated substrates for p97- dependent unfolding and subsequent proteasomal degradation or processing (Bays *et al*, 2001, Jarosch *et al*, 2002, Rape *et al*, 2001, Ye *et al*, 2001), whereas UBXN6 (also known as UBXD1) controls p97 functions in lysosomal and autophagic degradation pathways (Papadopoulos *et al*, 2017, Ritz *et al*, 2011) and SEP domain-containing cofactors mediate the maturation of protein phosphatase 1 complexes (Weith *et al*, 2018). At least in the case of p97-UFD1-NPL4, auxiliary cofactors from the UBA-UBX family can fine-tune the subcellular localization and/or substrate specificity of a major p97 complex (Buchberger *et al*, 2015, Schuberth & Buchberger, 2008). The yeast cofactor Ubx2 recruits Cdc48-Ufd1-Npl4 to the endoplasmic reticulum (ER) and mitochondria to promote ER– and outer mitochondrial membrane–associated protein degradation, respectively (Mårtensson *et al*, 2019, Neuber *et al*, 2005, Schuberth & Buchberger, 2005). Similarly, the metazoan cofactors UBXN7 and FAF1 were shown to direct p97- UFD1-NPL4 to nuclear chromatin to promote the removal of ubiquitylated substrates such as CDT1 and the CMG helicase (Franz *et al*, 2016b, Fujisawa *et al*, 2022, Kochenova *et al*, 2022, Tarcan *et al*, 2022).

The majority of cofactors interact with p97 through a small number of defined domains/motifs, including the UBX(-like) domain, the PUB and PUL domains, the SHP box, the VCP interacting motif (VIM) and the VCP binding motif (VBM) (reviewed in Buchberger *et al*, 2015). However, some p97 interactors do not possess any of these canonical binding motifs, suggesting that the current inventory of p97 cofactors is incomplete and, hence, that the full scope of the p97–cofactor network is far from being understood. One example illustrating this point is GIGYF1/2 (yeast Smy2), which was very recently shown to regulate p97 in the transcription stress response (Lehner *et al*, 2022).

Here, we report the identification of the previously uncharacterized, evolutionarily conserved FAM104 protein family as p97 cofactors. We show that FAM104 proteins bind p97 directly *via* a non-canonical helical motif and that they associate with p97-UFD1- NPL4 complexes in cells. We demonstrate that FAM104 proteins promote the nuclear localization of p97 and that their loss causes impaired growth and hypersensitivity to chemical inhibition of p97.

## Results

### FAM104 proteins are a conserved family of p97 interactors

In our ongoing efforts to identify p97 binding partners, we isolated multiple clones encoding the hitherto uncharacterized proteins FAM104A and FAM104B in a yeast two-hybrid screen of a human testis cDNA library. FAM104A and FAM104B are both expressed in various isoforms originating from alternative transcript variants (Fig. 1A). For FAM104A, the two-hybrid hits matched isoforms 1, 2, and 5, whereas isoforms 3 and 4, which share the N-terminal 73 residues with isoforms 1/2, were not isolated. For FAM104B, the two-hybrid hits matched isoforms 3 and 4, which differ by the insertion of a single valine residue after residue 40. Compared to these isoforms, isoforms 5 and 6 of FAM104B possess slightly different N termini, whereas isoforms 1 and 2 possess 30 divergent C-terminal residues and isoform 7 is truncated after 46 residues. Of note, those isoforms of FAM104A and B that were isolated in the two-hybrid screen possess significant sequence homology, suggesting that they are evolutionarily related. Indeed, FAM104-like proteins are present in vertebrates as well as in many invertebrates including insects, octopuses and echinodermata, whereas the occurrence of two distinct, FAM104A- and FAM104B-like homologs appears to be restricted to mammals (Fig. 1B, Suppl. Fig. S1A). Basically, FAM104 family members possess a predicted mono- or bipartite classical nuclear localization signal (cNLS) at or close to the N terminus and a C-terminal, highly conserved helical region, which are separated by largely unstructured stretches of variable length (Figs. 1AB, Suppl. Fig. S1B).

**Figure 1.**
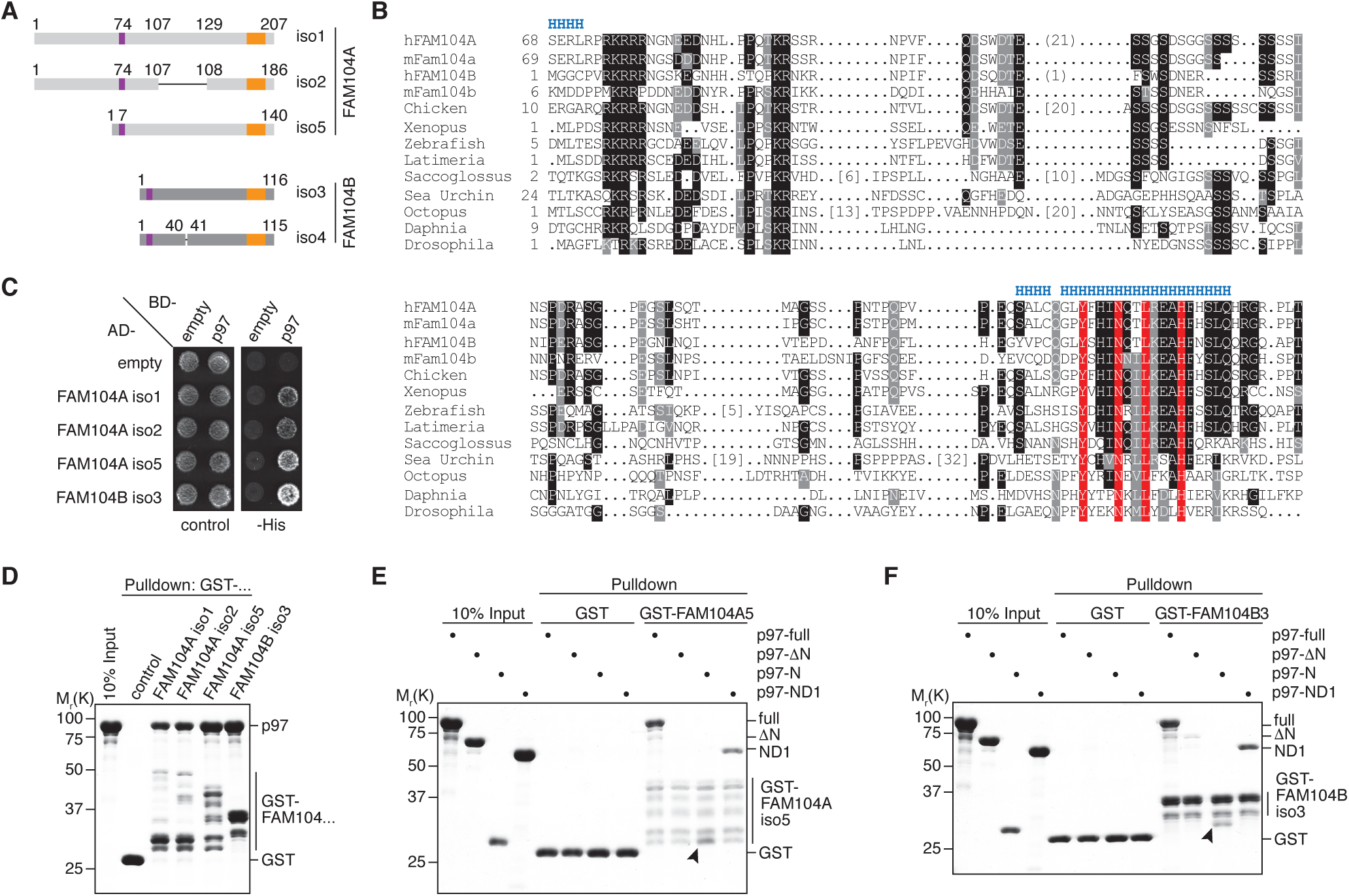
FAM104 proteins bind to p97. **(A)** Schematic overview of human FAM104A and FAM104B isoforms isolated in a yeast two-hybrid screen. Relevant amino acid residue numbers are shown, and conserved N- and C-terminal sequence motifs are indicated by purple and orange boxes, respectively. Internal deletions in FAM104A isoform 2 and FAM104B isoform 4 are indicated by thin lines. **(B)** Multiple sequence alignment showing representative members of the FAM104 family. Regions with predicted alpha-helical secondary structure are indicated at the top by ’H’. The most highly conserved residues in the C-terminal sequence motif are boxed in red. Other conserved residues are boxed in black or grey, according to the degree of conservation. Numbers in square brackets indicate the length of insertions. For human FAM104A and FAM104B, the sequences of isoforms 2 and 4, respectively, are shown, and numbers in round brackets indicate an 21-residue insertion present in isoforms 1 and 5 of FAM104A and a one-residue insertion present in isoform 3 of FAM104B, respectively. **(C)** Yeast two-hybrid analysis. Yeast PJ69-4a reporter cells transformed with the indicated combinations of bait (AD-) and prey (BD-) plasmids were spotted onto agar plates containing synthetic complete medium lacking uracil and leucine (control) or uracil, leucine, and histidine (-His). Growth was monitored after 3 days. **(D)** Glutathione sepharose pulldown assay using wild-type p97 and GST fusions of the indicated FAM104 proteins. Binding of p97 was analyzed by SDS–PAGE followed by Coomassie brilliant blue staining. **(E, F)** Glutathione sepharose pulldown assays as in (D), using the indicated p97 variants and GST fusions of FAM104A isoform 5 (E) and FAM104B isoform 3 (F), respectively. Arrowheads mark the position of the p97 N domain.

To validate the results of the two-hybrid screen, we tested isoforms 1, 2 and 5 of FAM104A and isoform 3 of FAM104B in directed yeast two-hybrid assays and confirmed that all four full-length proteins interact with p97 (Fig. 1C). Next, we performed glutathione sepharose pulldown experiments with purified GST fusions of these four FAM104 proteins and found that they were all able to efficiently bind recombinant p97 (Fig. 1D). In order to determine the FAM104A/B binding region of p97, we performed pulldown assays with FAM104A isoform 5 and FAM104B isoform 3, using different truncated variants of p97 (Figs. 1EF). The ND1 variant of p97 lacking the D2 ATPase domain as well as the N domain alone bound efficiently to both FAM104 proteins, whereas the ΔN variant lacking the N domain did not. Taken together, these results show that FAM104 proteins are a new family of evolutionarily conserved p97 interactors, and that the N domain is necessary and sufficient for the direct binding of p97 to FAM104A/B.

### FAM104 proteins bind to p97 *via* their C-terminal helix

As FAM104A/B do not possess any of the canonical p97 binding motifs found in other cofactors, we next sought to map their binding site for p97. The shortest FAM104A fragment isolated in the two-hybrid screen starts at glycine residue G131 of isoform 1 (equivalent to G110 and G64 of isoforms 2 and 5, respectively). Since the C-terminal alpha-helical region shows the highest sequence conservation among FAM104 family members (Fig. 1B) and is missing in those isoforms of FAM104A/B that were not isolated in the two-hybrid screen, we truncated this region from the C-terminus to test its importance for p97 binding (Fig. 2A). While deletion of the four C-terminal residues did not affect the two-hybrid interaction between FAM104A isoform 5 and p97, larger deletions of seven to 26 residues completely abolished p97 binding (Fig. 2B), even though the truncated proteins were expressed to similar levels in the reporter yeast strain (Fig. 2C). Consistent with this, deletion of the C-terminal 26 residues impaired the ability of all four FAM104A/B isoforms to bind p97 in a pulldown experiment (Fig. 2D), indicating that the C-terminal region contains the p97 binding site. We therefore tested if a peptide spanning the conserved alpha-helical and flanking residues (residues C180 to G203 in FAM104A isoform 1) can bind to p97. To that end, we immobilized the N-terminally biotinylated peptide on streptavidin sepharose beads and performed a pulldown assay with p97 (Fig. 2E). The peptide pulled down full-length p97 as well as the ND1 and N domain variants, demonstrating that the C-terminal conserved region of FAM104 family proteins is not only necessary, but also sufficient for p97 binding. We also noted some residual binding of the p97 ΔN variant, suggesting that the FAM104A- derived peptide has some weak affinity for a p97 region(s) outside the N domain.

**Figure 2.**
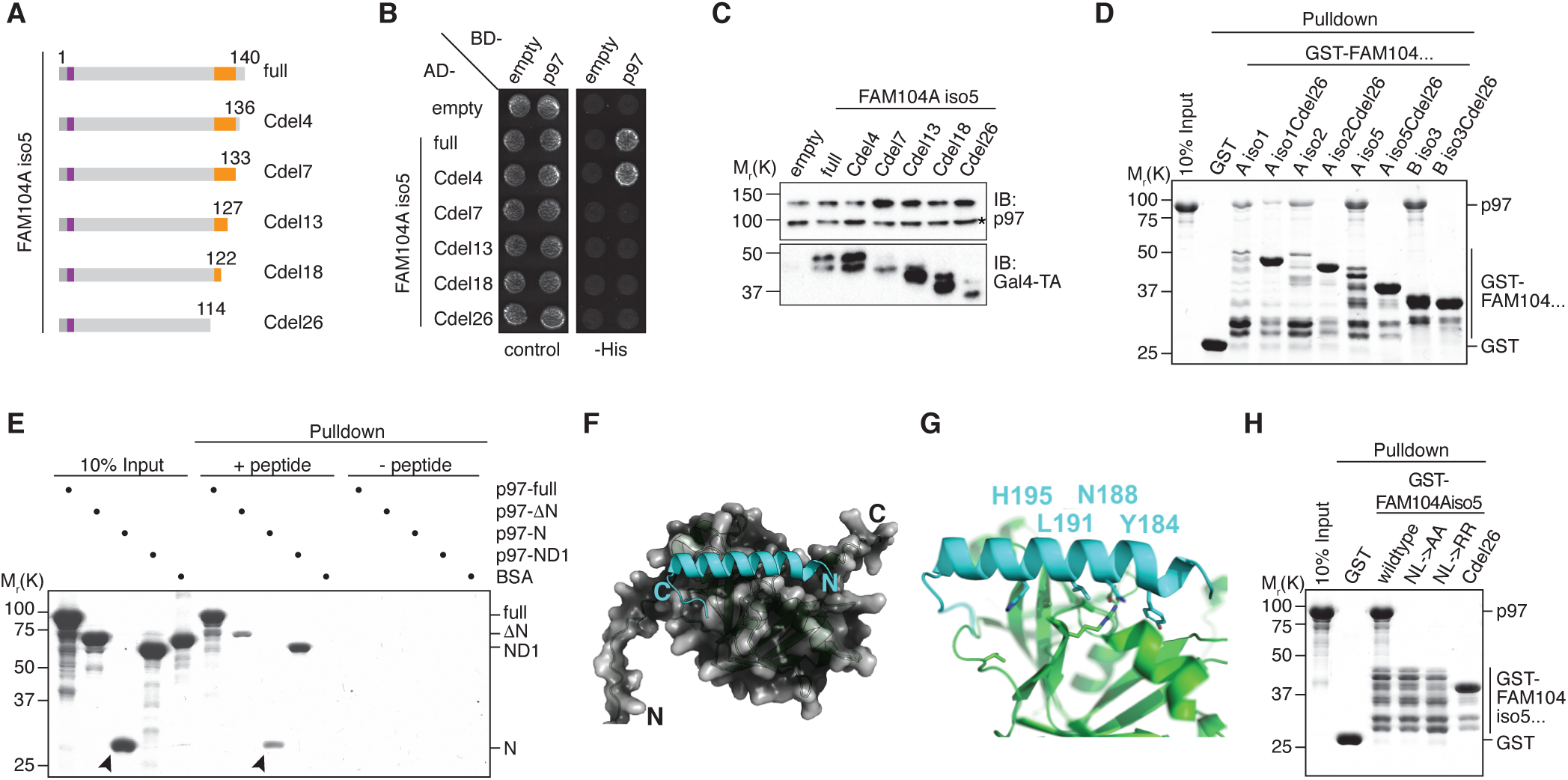
FAM104 proteins bind to p97 *via* their C-terminal helix. **(A)** Schematic overview of the C-terminal truncations of FAM104A isoform 5 used for yeast two-hybrid analysis. Labeling as in Fig. 1A. **(B)** Yeast two-hybrid analysis of p97 binding to the C-terminally truncated FAM104A isoform 5 variants shown in (A). **(C)** Expression levels of the two-hybrid fusion proteins in (B) were analyzed by Western blot (WB) using antibodies against p97 and the Gal4 transactivation domain (Gal4-TA). The asterisk in the p97 blot marks a cross-reactivity with endogenous Cdc48. **(D)** Glutathione sepharose pulldown assay using wild-type p97 and GST fusions of the indicated full- length or C-terminally truncated FAM104 proteins. **(E)** Streptavidin sepharose pulldown assay using the biotinylated peptide CQGLYFHINQTLREAHFHSLQHRG spanning the conserved C-terminal alpha-helix and flanking residues of FAM104A (residues C180 - G203 in isoform 1) and the indicated p97 variants. p97 binding to the immobilized peptide was analyzed by SDS-PAGE, followed by Coomassie brilliant blue staining. Arrowheads mark the position of the p97 N domain. **(F, G)** AlphaFold Multimer model of the C-terminal alpha-helix of FAM104A (turquoise) bound to the N domain of p97. (F) Overview showing binding to the subdomain cleft of the N domain. (G) Close-up view showing the interaction of the four most highly conserved residues with the N domain (green). Residue numbers refer to isoform 1 of FAM104A. **(H)** Glutathione sepharose pulldown assay using wild-type p97 and GST fusions of the indicated full-length (wildtype, NL->AA, NL->RR) or C-terminally truncated (Cdel26) variants of FAM104A isoform 5. NL->AA, N188A/L191A double mutant; NL->RR, N188R/L191R double mutant.

Using AlphaFold Multimer, we next generated a structural model of the C-terminal region of FAM104A bound to the N domain of p97 (Fig. 2FG). The central part of the C-terminal region was predicted as an alpha-helix that binds to the subdomain cleft of the p97 N domain (Fig. 2F), which is the major binding site for other N domain binding motifs including the UBX(-like) domain and the VIM and VBM linear motifs (Buchberger *et al*, 2015). Closer inspection of the modeled interface revealed that the four most strongly conserved residues of the helix, Y184, N188, L191 and H195, all contact the N domain (Fig. 2G; cf. Fig. 1B). The predicted interactions are centered around L191, which occupies a predominantly hydrophobic pocket in the p97 subdomain cleft.

Adjacent residues contribute to a hydrogen bonding network with p97, while Y184 is predicted to exhibit an orthogonal pi-stacking with Y138 of p97. To test the importance of this interface, we mutated the two central residues N188 and L191 to either alanine or arginine. We reasoned that the NL->AA double exchange should eliminate key contacts with the N domain, whereas the NL->RR double exchange should introduce steric clashes. Intriguingly, both double mutations abolished the interaction of FAM104A with p97 as efficiently as the deletion of the entire C-terminal region (Fig. 2H). Consistent with a conserved essential role of residues N188 and L191 for p97 binding, an AlphaFold Multimer model of the *Drosophila melanogaster* homologs of FAM104A/B (CG14229) and p97 (TER94) showed a highly similar binding interface with basically identical positions of the equivalent residues N90 and L93 (Suppl. Fig. S2). Together, our data identify the C-terminal alpha-helix of FAM104 proteins as a novel p97 binding motif and show that the strictly conserved residues N188 and L191 are of central importance for the interaction.

### FAM104 proteins bind to p97-UFD1-NPL4 in cells

To investigate if FAM104 proteins associates with p97 in cells, we ectopically expressed N-terminally FLAG epitope-tagged FAM104 proteins in HEK293T cells and performed anti-FLAG immunoprecipitations (Fig. 3A). Endogenous p97 was efficiently co-precipitated with all four wild-type FAM104 proteins tested, despite the fact that FAM104B isoform 3 was less well expressed than the three FAM104A isoforms and that the levels of soluble p97 in the input were strongly reduced upon ectopic expression of FAM104A isoforms 1 and 2. Importantly, UFD1 and NPL4 were co-precipitated at levels roughly proportional to those of p97, strongly suggesting that FAM104 proteins form complexes with p97-UFD1-NPL4. This conclusion is further supported by the finding that FAM104A Cdel26 variants lacking the C-terminal helix failed to co-precipitate not only p97, as expected, but also UFD1 and NPL4. (The expression level of the FAM104B isoform 3 Cdel26 variant was below the detection limit, precluding its analysis in this experiment.) Recently, p47, another major p97 cofactor, has been reported to interact with FAM104A (Raman *et al*, 2015). However, we were unable to detect a co-immunoprecipitation of FAM104 proteins and p47 under the same immunoprecipitation conditions that preserved the binding of FAM104A/B to p97-UFD1-NPL4 (Suppl. Fig. S3), even though our data do not exclude the possibility of a transient, conditional or weak interaction with p47.

**Figure 3.**
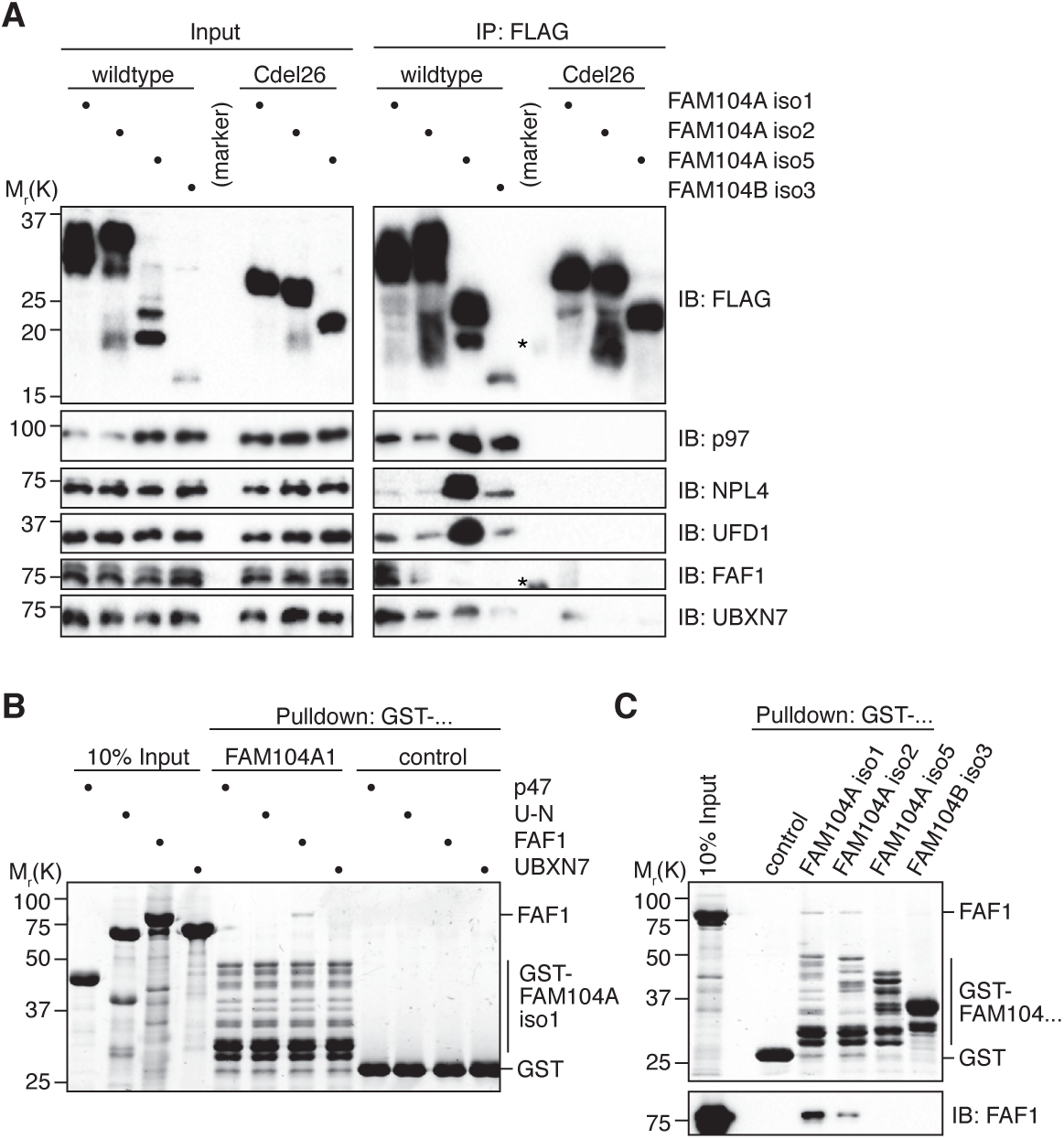
FAM104 proteins form complexes with p97, UFD1-NPL4 and UBA-UBX proteins in cells. **(A)** HEK293T cells ectopically expressing the indicated N-terminally FLAG-epitope- tagged wildtype or C-terminally truncated (Cdel26) FAM104 proteins were subjected to anti-FLAG immunoprecipitation (IP). Input and IP samples were immunoblotted for the FLAG epitope tag, p97 and the indicated p97 cofactors. The central empty lane had been loaded with a marker. The asterisks label marker bands cross-reactive with the FLAG and FAF1 antibody, respectively. **(B)** Glutathione sepharose pulldown assay, using the indicated p97 cofactors and GST-FAM104A isoform 1; U-N, UFD1-NPL4. **(C)** Glutathione sepharose pulldown assay, using FAF1 and GST fusions of the indicated FAM104 proteins. Upper panel, 2,2,2-Trichloroethanol-stained gel; lower panel, immunoblot with FAF1 antibody.

Interestingly, FAF1 and UBXN7, two auxiliary cofactors important for nuclear functions of p97-UFD1-NPL4, were also co-precipitated with the FAM104 proteins (Fig. 3A). Whereas FAF1 bound to isoforms 1 and 2 of FAM104A, UBXN7 bound to all four FAM104 proteins. Of note, the amounts of co-precipitated FAF1 and UBXN7 did not correlate with those of p97, UFD1 and NPL4, suggesting that the interaction of FAF1 and UBXN7 with FAM104A/B may not strictly depend on the p97-UFD1-NPL4 complex. To directly address this possibility, we performed pulldown experiments with purified p97 cofactors and found that FAF1, but not p47, UFD1-NPL4 or UBXN7 bound to FAM104A isoforms 1 and 2, whereas no binding of FAF1 to FAM104A isoform 5 and FAM104B isoform 3 was detected (Fig. 3BC). These results suggest that the N-terminal extension shared by FAM104A isoforms 1 and 2 mediates a direct interaction with FAF1.

Taken together, our data show that FAM104 proteins bind to p97-UFD1-NPL4 complexes in cells and that this binding is mediated by the C-terminal helix.

### FAM104 proteins increase nuclear p97 levels

All FAM104 proteins possess a *bona fide* cNLS at or close to the N terminus (Figs. 1AB, Suppl. Fig. S1B). To analyze their subcellular localization, we therefore ectopically expressed N-terminally FLAG epitope-tagged FAM104 proteins in HeLa cells and performed confocal immunofluorescence microscopy using anti-FLAG antibodies (Fig. 4A). Consistent with the very high score calculated by cNLS mapper (Kosugi *et al*, 2009), all four FAM104 proteins strongly accumulated in the nucleus. Intriguingly, endogenous p97 co-accumulated with FAM104A/B in the nuclei of transfected cells, as evident from the comparison with neighboring non-transfected cells or with the vector control (Fig. 4A). We next explored the effect of deleting the C-terminal helix or the cNLS of FAM104A, using C-terminally truncated isoforms 1 and 2 as well as cNLS- deleted isoforms 1, 2 and 5, respectively (Fig. 4B). The corresponding variants of the other FAM104A/B isoforms under study were poorly expressed below the detection limit of the immunofluorescence experiments, precluding their analysis (data not shown). The C-terminally truncated FAM104A Cdel26 variants localized to the nucleus, but failed to effectuate a significant nuclear accumulation of p97, as expected from the loss of their p97 binding site. The cNLS-deleted FAM104A variants showed a nuclear and cytoplasmic distribution, in accordance with their small size of less than 40 kDa, and a much less pronounced nuclear accumulation of p97 (see Suppl. Fig. S4A for higher intensity images of full-length *versus* cNLS-deleted FAM104A isoform 1). To quantify these results, we used CellProfiler (McQuin *et al*, 2018) for segmentation of the images into nuclei and cytoplasm (see Suppl. Fig. S4B for an example) and determined the ratio of nuclear to cytoplasmic p97 signal in control-transfected *versus* FAM104A isoform 1- transfected cells (Suppl. Fig. S4C). The ratio increased strongly from 2.03 ± 0.01 for control-transfected cells to 7.23 ± 0.54 for cells transfected with wild-type FAM104A isoform 1, reflecting the robust nuclear accumulation of p97 in the presence of FAM104A. By contrast, cells transfected with the cNLS-deleted variant showed only slightly elevated nuclear p97 levels (Suppl. Fig. S4AC). These differential effects on p97 localization correlated well with the nuclear levels of wild-type *versus* cNLS-deleted FAM104A isoform 1 (Suppl. Fig. S4AD). Together, these results suggest that FAM104 proteins target p97 *via* a piggy-back mechanism to the nucleus.

**Figure 4.**
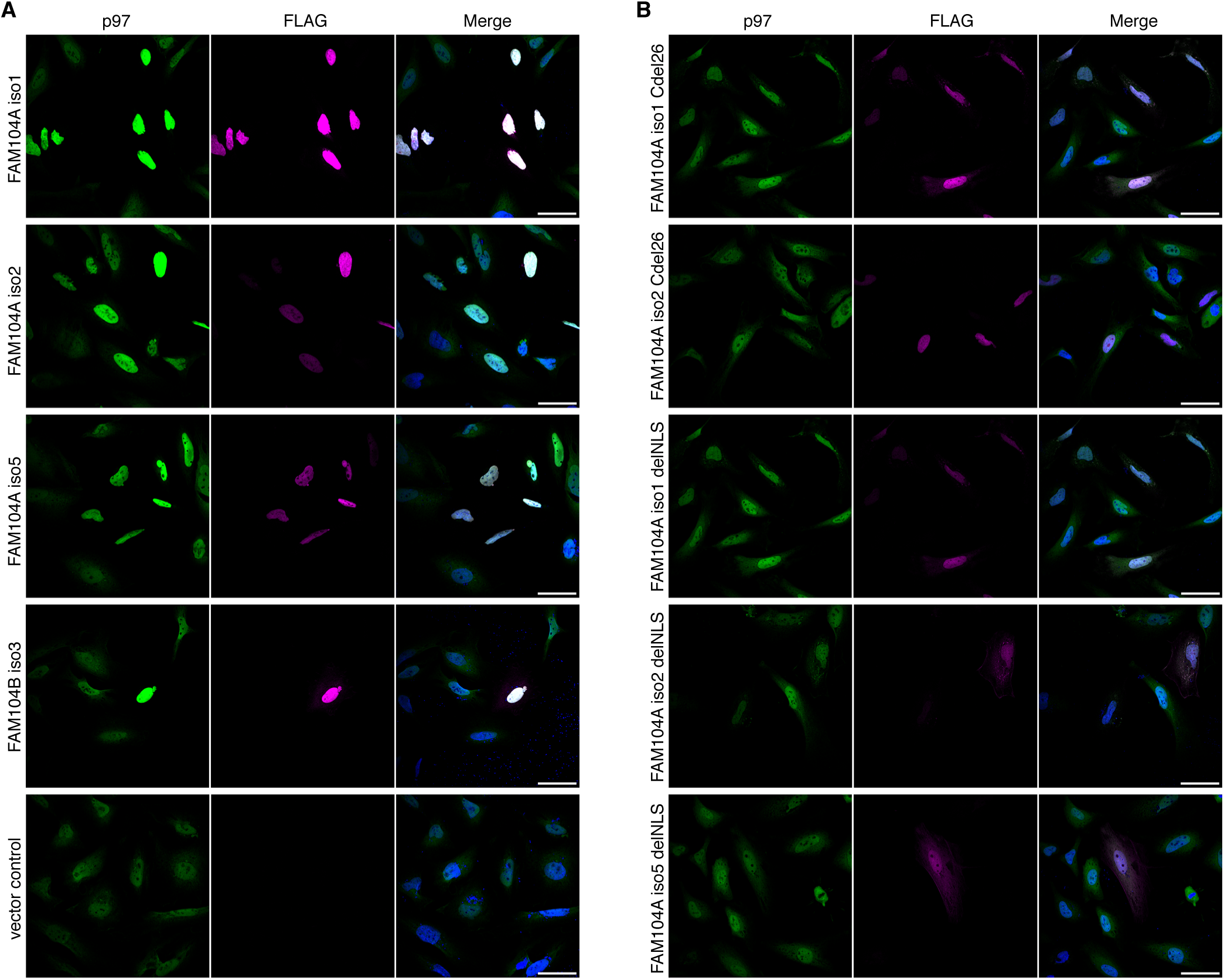
Ectopic expression of FAM104 proteins increases nuclear p97 levels. HeLa cells ectopically expressing the indicated, N-terminally FLAG epitope-tagged FAM104 proteins were analyzed by confocal immunofluorescence microscopy, using antibodies against endogenous p97 and the FLAG epitope. Scale bars, 50 µm. All images were taken with identical acquisition settings and processed identically. **(A)** HeLa cells expressing full-length FAM104A isoforms 1, 2 and 5, FAM104B isoform 3, and empty vector control. **(B)** HeLa cells expressing FAM104A isoforms 1 and 2 lacking the C-terminal conserved helix (Cdel26) or the cNLS (delNLS), and FAM104A isoform 5 lacking the cNLS.

As the overexpression of FAM104A isoforms 1 and 2 had caused a noticeable reduction in the p97 input levels in the immunoprecipitation experiments described above (Fig. 3A, Suppl. Fig. S3), we hypothesized that a subpopulation of p97 may have become insoluble under these conditions. To test this possibility, we performed a biochemical fractionation of HEK293T cells ectopically expressing FAM104A isoform 1 or 2 (Fig. 5). Both isoforms strongly accumulated in the soluble nucleoplasmic and insoluble chromatin fractions and were only weakly detected in the cytoplasmic fraction, in full agreement with the immunofluorescence experiments. Intriguingly, the overexpression of both FAM104A isoforms did not affect the total amount of p97, as judged by the signal in SDS-denatured total cell extracts, but caused a strong increase of p97 in the chromatin fraction at the expense of the cytoplasmic pool. Because the other two FAM104A/B isoforms under study were only weakly expressed, their potential to cause similar effects could not be addressed (data not shown). Our data thus indicate that FAM104A isoforms 1 and 2 specifically promote the chromatin association of p97, and they provide a rationale for the reduction of soluble p97 observed in the immunoprecipitation experiments.

**Figure 5.**
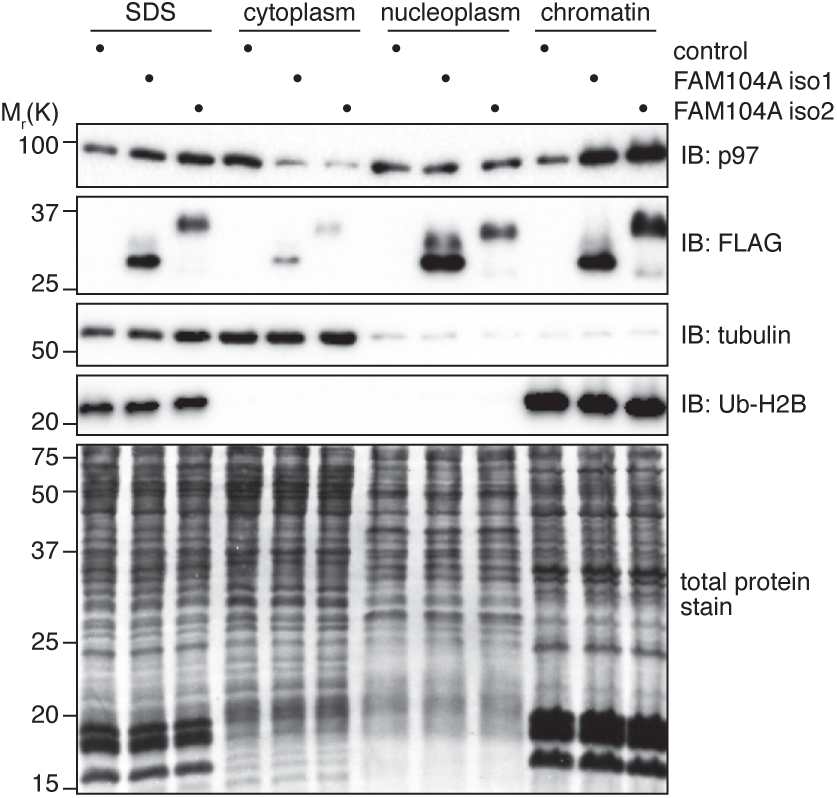
Ectopic expression of FAM104 proteins promotes the association of p97 with chromatin. HEK293T cells were transfected with plasmids encoding the indicated N-terminally FLAG-epitope-tagged FAM104 proteins or with empty vector (control). Total protein extracts were prepared by direct boiling part of the cells in SDS PAGE sample buffer (SDS). The remaining cells were processed to cytoplasmic, soluble nuclear (nucleoplasmic) and chromatin fractions as indicated. Tubulin and ubiquitylated histone H2B (Ub-H2B) served as markers for the cytoplasmic and chromatin fractions, respectively, whereas the Coomassie staining of the membrane (total protein stain) served as loading control.

Since the previous experiments relied on the ectopic expression of FAM104 proteins, we next sought to determine if endogenous FAM104A/B also has an impact on the nuclear localization of p97. To this end, we generated FAM104A knockout as well as FAM104A+B double knockout HeLa cell pools by CRISPR/Cas9-mediated genome editing. Compared to control cells, both knockout cell pools showed a reduced nuclear p97 signal, with the ratio of nuclear to cytoplasmic p97 decreased by about 22% (Figs. 6AC). The siRNA-mediated depletion of FAM104A had a similar effect (Figs. 6BD), which was more obvious to the naked eye in maximum intensity projections of z stacks (Suppl. Fig. S4E). Moreover, a comparable reduction in nuclear p97 was observed in FAM104 single and double knockout HEK293T cell pools (Suppl. Fig. S4F). In summary, these data demonstrate that endogenous FAM104 proteins modulate the nucleo-cytoplasmic distribution of p97.

**Figure 6.**
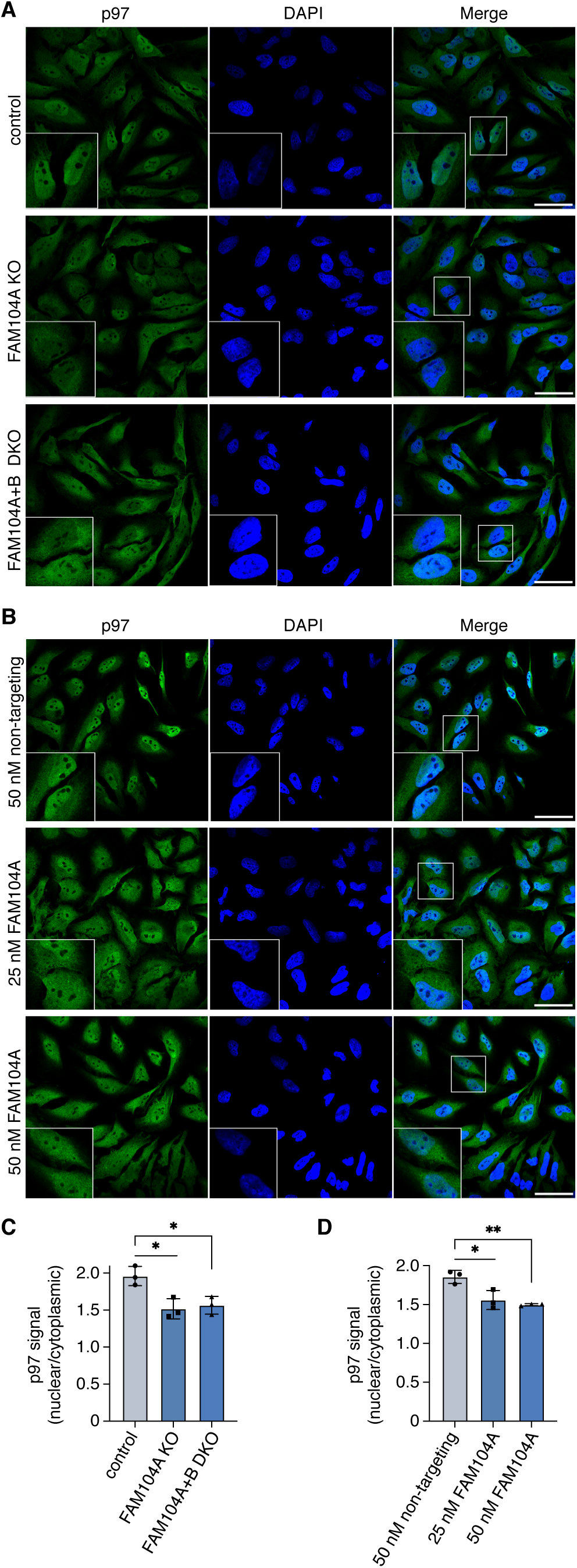
Depletion of FAM104 proteins reduces nuclear p97 levels. **(A)** Control, FAM104A single knockout (KO) and FAM104A/FAM104B double knockout (DKO) HeLa cell pools were analyzed by confocal immunofluorescence microscopy, using an antibody against endogenous p97. **(B)** HeLa cells were transfected with non-targeting or FAM104A-specific siRNAs at the indicated final concentrations for 72 hours and analyzed as in (A). **(C, D)** Quantification of the ratios of nuclear to cytoplasmic p97 signals in (A) and (B), respectively. Shown is the mean ± s.d.; n = 3 with ≥ 80 cells per replicate and condition; *, p <0.05; **, p < 0.01; Student’s t-test. Scale bars, 50 µm.

### FAM104 proteins promote normal cell growth

p97 functions in a number of important nuclear processes. To analyze if the reduction of nuclear p97 in the absence of FAM104 proteins has consequences for the cell fitness, we performed growth assays with HeLa FAM104A/B double knockout cells (Fig. 7). Eight days after seeding equal numbers of cells, the number of knockout cells was reduced by more than 40% compared to control cells, indicating that FAM104 proteins are required for normal growth under otherwise unperturbed conditions (Fig. 7A). Next, we determined the growth of FAM104 knockout cells under conditions of limited p97 activity, i.e. in the permanent presence of low, sub-lethal concentrations of the ATP-competitive p97 inhibitor CB-5083 (Anderson *et al*, 2015), for seven days. While no statistically significant growth difference between knockout and control cells was observed at lower CB-5083 concentrations, growth of the knockout cells was significantly impaired in the presence of 100 and 200 nM CB-5083, indicating that loss of FAM104A/B made the cells hypersensitive to p97 inhibition (Fig. 7B). Note that growth of both cell lines was normalized to their respective growth in the absence of CB-5083, thus compensating for the slower growth of the knockout cells under unperturbed conditions. Finally, we analyzed the growth of FAM104A/B knockout cells in the presence and absence of CB-5083 after pre-treatment with the DNA damage-inducing topoisomerase I inhibitor, camptothecin (Fig. 7C). In the absence of CB-5083, the relative growth of camptothecin-treated control and knockout cells was comparable, indicating that loss of FAM104A/B alone does not cause a hypersensitivity to camptothecin. By contrast, a clear trend towards reduced growth was observed for the knockout cells in the presence of CB-5083. While this effect did not reach statistical significance at all CB- 5083 concentrations, it did so at 25 nM, i.e. at a concentration that had no effect in the absence of camptothecin pretreatment (see Fig. 7B). Together, these results suggest that FAM104 proteins are not only required for normal cell growth, but also for normal DNA damage repair under conditions of limited p97 activity.

**Fig. 7:**
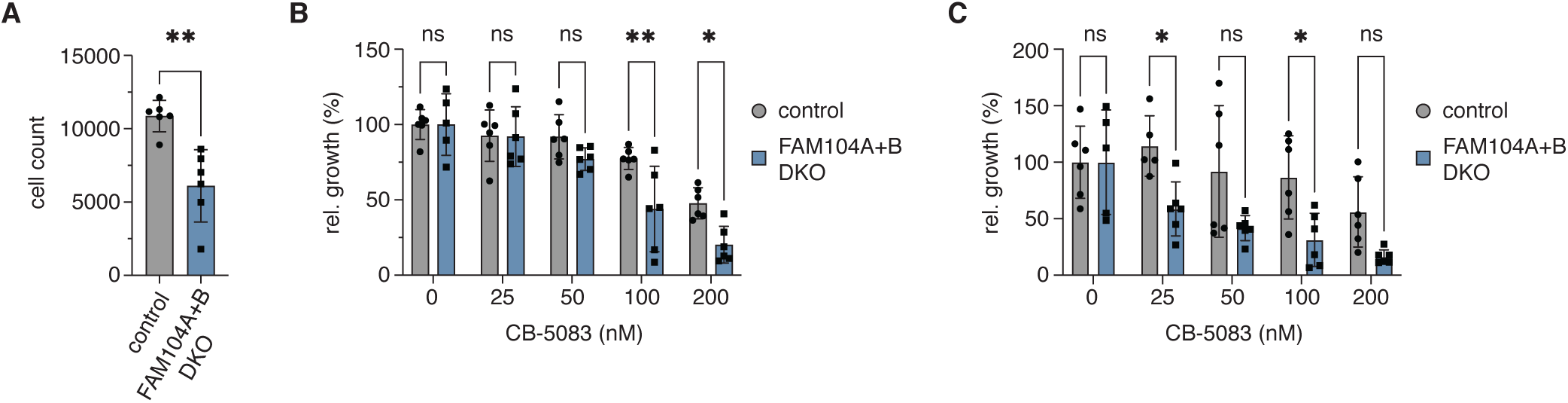
Deletion of FAM104A/B causes reduced growth and hypersensitivity to p97 inhibition. HeLa FAM104A/B double knockout (DKO) and control cells were seeded in 96 well plates, cultivated in the absence and presence of CB-5083, stained with Hoechst dye and automatically counted using a high-content microscopy platform. n=6, shown are mean ± s.d.; *, p <0.05; **, p < 0.01; ns, not significant. **(A)** Absolute cell count of untreated cells after 8 days. A Student’s t-test was performed. **(B)** Relative growth at the indicated concentrations of the p97 inhibitor CB-5083. Values were normalized separately for control and knockout cells to the respective mean value at 0 nM CB-5083. A two-way ANOVA was performed using the normalized data. **(C)** Same as in (B), but cells were pre-treated with 1 uM camptothecin for 70 min before starting the CB-5083 treatment.

## Discussion

In this study, we characterized FAM104 proteins as a family of evolutionarily conserved p97 interactors and identified their C-terminal helix as a novel p97 binding motif. We showed that FAM104 proteins associate with p97 complexes containing the cofactors UFD-NPL4, FAF1 and UBXN7 in living cells, and that they recruit p97 to the nucleus and nuclear chromatin. Loss of FAM104 proteins reduced nuclear p97 levels, impaired cell growth and caused hypersensitivity to p97 inhibition in the absence and presence of DNA damage, indicating that FAM104 proteins support optimal p97 activity in the nuclear compartment.

Whereas FAM104B had not been reported to interact with p97 before, FAM104A was identified as a p97 interactor in several high-throughput proteomic and yeast two-hybrid screening projects (Fogeron *et al*, 2013, Haenig *et al*, 2020, Hein *et al*, 2015, Huttlin *et al*, 2017, Rolland *et al*, 2014). However, the interaction between p97 and FAM104A was not further validated in any of these previous studies. In fact, the validation of p97 binding is complicated by the existence of five different isoforms, of which only three (isoforms 1, 2 and 5) contain the p97-interacting C-terminal helix. The existence of p97 binding and non-binding isoforms of FAM104A in humans also initially obscured the prevalence of FAM104 proteins in the animal kingdom. Nevertheless, bioinformatic searches using the C-terminal p97 binding helix as query readily identified homologs in all five vertebrate classes as well as in *Saccoglossus* and sea urchin, leading to the subsequent identification of numerous invertebrate homologs (Fig. S1A). Outside of the chordates, FAM104 proteins are found in many arthropods (insects, spiders, crustaceans) and selected other invertebrate lineages including mollusks (octopi, snails), tardigrades and rotifers. FAM104 homologs are absent in nematodes, protists, plants and fungi. The three human p97-interacting FAM104A isoforms possess an N-terminal 74-residue extension exclusively found in eutheria, a 21-residue insertion after residue 107 (numbering referring to isoform 1/2) that is found in mammals, birds and reptiles, or both (Figs. 1AB). Amphibian, fish and invertebrate homologs contain neither insertion and are therefore more similar to human FAM104B with respect to their overall organization. The widespread occurrence of FAM104 proteins strongly suggests a conserved function, even though very little information is available for non-human homologs. The fly homolog CG14229 lacks any functional annotations, whereas mouse Fam104a was reported to interact with ubiquitin (Zhang *et al*, 2017). The binding preference for K6-, K48- and K29-linked di-ubiquitin observed in that study closely resembled that of p97, suggesting that Fam104a does not bind ubiquitin directly, but in the context of a p97–cofactor complex such as p97-Ufd1-Npl4.

Here, we showed for the first time that FAM104 proteins and p97 interact directly *via* their highly conserved C-terminal helix and N domain, respectively (Figs. 1 & 2). Structural modeling revealed that the C-terminal helix binds to the subdomain cleft of the p97 N domain. Interestingly, the position of the FAM104A helix in the model is similar to those of the helical VIM and VBM p97 binding motifs in the absence of any significant sequence similarity (Buchberger *et al*, 2015, Hanzelmann & Schindelin, 2011, Lim *et al*, 2016). Even the respective key p97-contacting residues are distinct, with N188 and L191 in the C-terminal helix of FAM104 *versus* predominantly arginine residues in VIM and VBM. The finding that alpha-helices can efficiently bind to the N subdomain cleft *via* just one or two key residues underlines the plasticity of this binding interface and raises the intriguing possibility that additional interactors bind p97 through yet other helical motifs.

Our cell-based experiments revealed a significant effect of FAM104 proteins on the subcellular localization of p97 (Figs. 4-6). The overexpression of FAM104 proteins resulted in a strong increase in p97 nuclear localization and chromatin association, whereas both CRISPR/Cas9-mediated deletion and siRNA-mediated depletion caused a decrease in nuclear p97 levels. While we were unable to detect endogenous FAM104 proteins due to the lack of suitably sensitive antibodies, the highly consistent effects of the deletion and depletion experiments on nuclear p97 levels provide strong indirect evidence that endogenous FAM104 proteins indeed localize to the nucleus, in agreement with their potent cNLS and the robust nuclear accumulation of ectopically expressed FAM104 proteins. Because the de(p)letion of FAM104A alone had similar effects as the double knockout of FAM104A and FAM104B in these experiments, FAM104A apparently plays a more important role for the nuclear localization of p97 than FAM104B, at least in HeLa and HEK293T cells. Interestingly, the bi-partite cNLS found in yeast Cdc48 is not conserved in human p97 (Madeo *et al*, 1998), which appears to possess a relatively weak potential cNLS (residues K60-R65, score 5 according to cNLS mapper (Kosugi *et al*, 2009). However, available three-dimensional structures of p97 suggest that this potential cNLS is part of a beta-hairpin in the N domain that is not constitutively exposed due to nucleotide-dependent conformational changes and cofactor binding. We therefore propose that FAM104 proteins promote the nuclear import of p97 by exposing a very strong cNLS that is connected *via* a long, flexible region with the C-terminal helix tightly interacting with the p97 N domain. However, since de(p)letion of FAM104 proteins caused only a partial reduction in nuclear p97 levels, it is likely that additional nuclear cofactors such as UBXN7 contribute to the efficient nuclear targeting of p97.

We were able to show that FAM104 proteins associate with the major p97-UFD1- NPL4 complex (Fig. 3), potentially implicating FAM104 proteins in nuclear p97 functions such as DNA damage repair, regulation of DNA replication, and control of transcription. Consistent with this possibility, isoforms 1 and 2 of FAM104A bound the auxiliary cofactors UBXN7 and FAF1 (Fig. 3) and promoted the chromatin association of p97 (Fig. 5). While the finding that FAM104A/B knockout cells are hypersensitive to p97 inhibition in the absence and presence of the DNA damaging agent camptothecin (Fig. 7BC) is in line with an involvement of FAM104 proteins in p97-dependent DNA damage repair processes, we were unable to detect a camptothecin sensitivity in the absence of p97 inhibitor. It is possible that the reduction in nuclear p97 levels by 22% upon FAM104A/B knockout is insufficient to severely compromise p97-dependent nuclear processes, and/or that these essential processes are safeguarded by redundant p97 cofactors.

The nuclear recruitment function of FAM104 proteins is reminiscent of two other small p97 interactors, SVIP and VIMP (also known as SELENOS, selenoprotein S). SVIP is myristoyl-anchored to the lysosomal membrane, whereas VIMP is a single-spanning ER membrane protein. It therefore appears that FAM104 proteins, SVIP and VIMP recruit p97 to distinct subcellular localizations with high demand for p97 activity.

According to the Human Protein Atlas and Proteomics DB databases, FAM104A and B are broadly expressed in many human tissues, including strong expression in the brain. In this context, it is interesting to note that both FAM104 proteins have recently been implicated in neurodegenerative diseases (FAM104A: Amyotrophic lateral sclerosis; FAM104B: Alzheimer’s disease), based on systematic yeast two-hybrid screening and computationally predicted neurodegenerative disease-associated protein clusters (Haenig *et al*, 2020). Moreover, the potential link between FAM104A/B and p97- mediated DNA damage repair suggested by our data could hint to a role of FAM104A/B in p97-dependent cancer cell survival, an area of intense biomedical research efforts.

While the exact role of FAM104 proteins in the control of nuclear p97 functions remains to be determined, this work provides a starting point for the future exploration of the FAM104 protein family in basic and translational research.

## Methods

Plasmids, antibodies and key chemicals used in this study are listed in Supplementary Table S1.

### Plasmids

Full-length and C-terminally truncated coding sequences of human FAM104A isoforms 1 (NCBI reference sequence NM_001098832.2), 2 (NM_032837.3) and 5 (NM_001289412.2) and of human FAM104B isoform 3 (NM_001166700.2) were PCR-amplified from a yeast two-hybrid human testis cDNA library (Clontech, cat. no. 638810) and cloned *via* appropriate restriction sites into pGAD-C1 (James *et al*, 1996), pGEX-4T1 (Cytiva) and pCMV-Tag2B (Agilent) using standard procedures. The coding sequence of FAM104A isoform 5 was mutated using the QuikChange XL II kit (Agilent) with primers FAM104A_NL-RR_fwd (5’-cct cta ctt cca cat ccg cca gac ccg cag gga ggc cca ctt cc), FAM104A_NL-RR_rev (5’- gga agt ggg cct ccc tgc ggg tct ggc gga tgt gga agt aga gg), FAM104A_NL-AA_fwd (5’- cct cta ctt cca cat cgc cca gac cgc gag gga ggc cca ctt cc) and FAM104A_NL-AA_rev (5’- gga agt ggg cct ccc tcg cgg tct ggg cga tgt gga agt aga gg) according to the manufactureŕs instructions. The cNLS-deleted variants of FAM104A isoforms 1 and 2 (deletion of codons 74-82) and FAM104 isoform 5 (deletion of codons 2-15) were generated by two- and one-step PCR reactions, respectively. The coding sequence of human p97 was cloned into pGBDU-C1 (James *et al*, 1996). The coding sequence of human UBXN7 was cloned into mini-pRSETA (Perrett *et al*, 1999). The identities of all inserts were confirmed by Sanger sequencing. Plasmids for the bacterial expression of FAF1 (Jensen *et al*, 2001), p47 (Allen *et al*, 2006), UFD1-NPL4 and full-length and truncated p97 (Fernandez-Saiz & Buchberger, 2010, Rothballer *et al*, 2007) have been described.

### Proteins and peptides

Expression of GST-tagged FAM104A/B proteins or of hexahistidine-tagged UBXN7 in *E. coli* BL21(DE3)pRIL (Agilent) was induced with 1 mM IPTG during overnight growth at 18 °C. GST fusion proteins were affinity-purified on a glutathione sepharose matrix (Cytiva) according to the manufacturer’s instructions. Purified GST fusion proteins were dialyzed against TBS (50 mM Tris-HCl, 150 mM NaCl, pH 7.5) containing 2 mM DTT, aliquoted, flash frozen in liquid nitrogen and stored at -80 °C. His_6_-UBXN7 was purified by Ni^2+-^NTA affinity chromatography (Qiagen) according to the manufacturer’s instructions, followed by anion exchange chromatography on a ResourceQ column (Cytiva). Purified His_6_-UBXN7 was dialyzed against 50 mM Tris-HCl, 50 mM NaCl, 2 mM DTT, pH 8.0 and flash frozen. His_6_-FAF1, His_6_-p47 (Allen *et al*, 2006) and UFD1- NPL4, full-length and truncated p97 (Fernandez-Saiz & Buchberger, 2010) were purified as previously described.

### *In vitro* binding assays

Pulldown assays using immobilized GST fusion proteins or biotinylated peptide (Biotin- CQGLYFHINQTLREAHFHSLQHRG-COOH; PANATecs GmbH, Tübingen, Germany) were essentially performed as described (Böhm *et al*, 2011), using 10 µl glutathione or streptavidin sepharose beads, respectively, 0.76 nmole GST fusion or 5 nmole peptide, respectively, and 0.2 nmole (monomer) p97 or 0.5 nmole His_6_-FAF1 in pulldown buffer (TBS containing 2 mM DTT and 1% or 0.1% Triton X-100 for glutathione and streptavidin pulldowns, respectively).

### Bioinformatic analyses

Sequence alignments were generated using the L-INS-I algorithm of the MAFFT package (Katoh & Standley, 2013). The multiple alignments were used for the generation of generalized profiles using pftools (Bucher *et al*, 1996). Generalized profile searches were performed using all proteins from the Uniprot database (https://www.uniprot.org). Protein clustering was performed using the CLANS software (Frickey & Lupas, 2004). Sequence logos were generated using the WebLogo server (https://weblogo.berkeley.edu)

### Structural modeling

For structure prediction, the sequence of the p97 N domain (residues 1-213 of human p97) and the sequence of the FAM104A C-terminal region (residues 180-207 of human FAM104A isoform 1) were fed into a local installation of AlphaFold Multimer running locally with two GPUs (Evans *et al*, 2022). Five models were created, which showed no visually discernible differences for the interface in the central region, accompanied by a high AlphaFold confidence score around residue L191 of FAM104A. A similar approach was used for the *Drosophila melanogaster* homologs, which created highly similar results.

### Mammalian cell culture

HeLa (ATCC; CCL-2) and HEK293T (ATCC; CRL-3216) cells and their derivatives were cultured in Dulbecco’s modified Eagle’s medium (DMEM) supplemented with 10% fetal bovine serum and 1% penicillin/ streptomycin in a humidified atmosphere with 5% CO_2_ at 37 °C. In addition, 1.5 μg/ml puromycin was added to the culture media for HeLa and HEK293T knockout cell pools.

To create *FAM104A/B* knockout cell pools, recombinant lentiviruses were produced using pLentiCRISPRv2-derived plasmids encoding gRNAs targeting the human *FAM104A* and *FAM104B* genes, respectively, generated using 3Cs technology (Wegner *et al*, 2019) (gift from Manuel Kaulich, University of Frankfurt). The following gRNA sequences were used:

Fam104A1-KO-2-R_79_CGTAGCTTCCATCCGCCAGC;

Fam104A1-KO-3-R_38_CCTCGGGCCTTGGCTCTCGC;

Fam104A2-KO-1-R_190_TGTCCGGGCTATTGATGCTG;

Fam104A2-KO-2-R_170_ACCGCGCAGAACGCTTTGTT;

Fam104A3-KO-1-R_172_CTCCGCGAAGAGAGGGAACA;

Fam104A3-KO-3-R_70_CCGAAACACAACCCCCTCTG;

Fam104B1-KO-1-R_187_CTGTATCTTGAGAATCCTGA;

Fam104B1-KO-2-R_18_CTTGCTCTCTCTGGGATATT;

Fam104B1-KO-3-R_139_CATTAATATCCCAGAGAGAG;

Fam104B2-KO-1-R_19_GATTGTTACTGAACCCGATG;

Fam104B2-KO-2-R_85_GTTTTCATGGAGTGATAATG;

Non-Human-Target-309-KO-1-R_156, AACATGACGTTCAAGATTGG;

Non-Human-Target-365-KO-5-R_5, ACCACTGTTCTACGCGCAGG;

Non-Human-Target-415-KO-2-R_24, TTGAACGGGCCGCGGAAGCG;

Non-Human-Target-42-KO-15-R_115, CCCGCATGACACCGTCACTT.

After production of recombinant lentiviruses in HEK293T cells by co-transfection of the pLentiCRISPRv2 constructs with pMD2.G and psPAX2 (Addgene plasmids #12259 and #12260; gifts from Didier Trono), HeLa and HEK293T cells at 40 - 60% confluence were transduced with lentivirus-containing supernatant mixed with polybrene to a final concentration of 8 μg/ml. 48 h after transduction, fresh culture medium containing 1.5 µg/ml puromycin was added, and the cell pools were kept under constant selection.

For the ectopic expression of FAM104A/B, cells were transfected at 60% confluence using polyethylenimine and analyzed 42 h after transfection. For siRNA-mediated depletion of FAM104A, cells were seeded in 12-well plates and transfected with FAM104 or control siRNA (25 or 50 nM final concentration) using 1.2 ul Oligofectamine (Thermo Fisher) diluted in 100 ul Opti-MEM (Thermo Fisher). After 20 h, the medium was changed and the knockdown was continued for a total of 72 h.

### Immunoprecipitation experiments

For immunoprecipitation of FLAG-epitope-tagged FAM104 proteins, HEK293T cells were transfected with pCMV-Tag2B encoding the indicated FAM104A/B variants or with empty vector. 48 h after transfection, cells were harvested, washed twice with cold PBS (140 mM NaCl, 2.7 mM KCl, 10 mM Na_2_HPO_4_, 1.8 mM KH_2_PO_4_) and resuspended in 1000 μl of lysis buffer (50 mM Tris-HCl pH 7.6, 150 mM NaCl, 2 mM MgCl_2_, 1% NP40, 10% glycerol, 1 mM DTT) containing protease inhibitors (1 mM PMSF, 1x Roche complete protease inhibitor mix). The cells were lysed on ice for 10 min, and lysates were sonicated six times for 9 sec in a Branson Sonifier using a microtip at 35% output control. cell debris was removed by centrifugation (20,000 g, 20 min, 4 °C), and the protein concentration of the supernatants was determined with the Pierce BCA Protein Assay Kit (Thermo Fisher) and adjusted to equal input levels. 50 ul were taken as input sample, mixed with five-fold concentrated SDS PAGE sample buffer (250 mM Tris-HCl pH 6.8, 10% (w/v) SDS, 30% glycerol, bromophenolblue) and heat-denatured. 2 mg of the input were incubated overnight at 4 °C with 50 μl FLAG M2 agarose beads on a rotating wheel. Beads were collected by centrifugation (1,400 g, 4 min, 4 °C), and the supernatant was aspirated. The beads were washed twice with lysis buffer containing protease inhibitors, once with lysis buffer without inhibitors, and once with TBS (50 mM Tris-Cl, pH 7.5, 150 mM NaCl). The immunoprecipitates were heat-denatured on the beads in 1x SDS PAGE sample buffer and further analyzed by immunoblotting.

### Immunoblotting

Protein samples were separated by SDS-PAGE and transferred onto PVDF membrane (Millipore) by semi-dry blotting using 1x Tris-glycine buffer (192 mM glycine, 25 mM Tris base, pH 8.3) supplemented with 20% methanol. The membrane was blocked with 5% milk in TBST (50 mM Tris-HCl pH 7.5, 150 mM NaCl, 0.1% Tween 20) and incubated with the indicated primary antibody in blocking solution overnight at 4°C. The membrane was washed with TBST (3x 10 min), incubated with HRP-conjugated secondary antibody (Dianova) diluted 1:7,500 in blocking solution for 1 h at RT, washed again with TBST (3x 10 min), and incubated with Clarity Western ECL Substrate (Bio-Rad). Chemiluminescence signals were detected using the Gel Doc XR+ system (Bio-Rad), and immunoblot images were processed with Image Lab (Bio-Rad).

### Immunofluorescence

Cells grown on coverslips to 60% confluence were washed twice with PBS, fixed using 3.7% formaldehyde in PBS for 12 min at RT, washed twice with cold PBS, permeabilized with 0.2% Triton X-100 in PBS for 10 min at RT, washed with PBS, and blocked by incubation with 2% BSA in PBS for one hour at RT. Cells were incubated with the indicated primary antibodies (diluted in 2% BSA in PBS) overnight at 4 °C, washed for 15 min with PBS and incubated with the appropriate fluorophore-coupled secondary antibodies for 2 h at RT. Hoechst staining was performed by incubating the cells for 10 min in PBS containing 2.5 μg/ml Hoechst 33342. Following this, cells were washed for 15 min with PBS and rinsed with water. Coverslips were mounted for microscopy with ProLong™ Glass Antifade Mountant and sealed with nail polish.

### Microscopy and image processing

Confocal immunofluorescence microscopy was performed at the Imaging Core Facility (Biocenter, University of Würzburg) using a Leica TCS SP2 confocal microscope equipped with an acousto optical beam splitter. Images were acquired using a 63x/1.4 oil immersion objective and Leica confocal software. Where higher resolution was required, 2x digital zooming was applied. Single planes or Z stacks were acquired using Diode UV (405 nm), Ar (488 nm), and HeNe (561 nm) lasers with 3 PMTs set to 419–452 nm, 497– 541 nm, and 595-650 nm, respectively. Image processing was performed using Fiji (Schindelin *et al*, 2012) and CellProfiler (McQuin *et al*, 2018). In CellProfiler, a pipeline was created to define nuclei *via* the Hoechst staining (thresholding method: minimum cross-entropy) and cells *via* the p97 staining (propagation from nuclei, thresholding method: otsu). The cytoplasm was defined by subtracting nuclei from cells. Following this segmentation, intensities were measured in the nucleus and the cytoplasm, and their ratio was calculated.

### Chromatin fractionation

To perform cellular fractionations, HEK293T cells were seeded in 10 cm dishes and transfected with pCMV-Tag2B encoding the indicated FAM104A isoforms or with empty vector. 24 h post transfection, cells were washed twice with cold PBS, harvested using cell scrapers and centrifuged for 5 min at 1.300 rpm. 10% of the cell material was centrifuged separately, directly resuspended in 100 μl SDS PAGE sample buffer and heat-denatured. The remaining cell pellets (containing around 3*10^7^ cells) were resuspended in 600 μl hypotonic buffer (20 mM Hepes-KOH pH 8.0, 5 mM KCl, 1.5 mM MgCl_2_, 1 mM DTT), incubated on ice for 20 min and lysed using a 27 gauge needle (10 strokes). The cytoplasmatic fraction was separated by centrifugation (10 min, 16.000 rpm, 4°C) from the nuclei-containing pellet and transferred to a separate reaction tube. After protein concentration determination and volume adjustment, 50 μl were taken, mixed with five-fold concentrated SDS PAGE sample buffer and heat-denatured. The pellet was resuspended in 400 μl nuclear extraction buffer (15 mM Tris-HCl pH 7.5, 1mM EDTA, 0.4 M NaCl, 10% sucrose, 1 mM DTT) and incubated on ice for 30 min with occasional vortexing. The chromatin-containing insoluble fraction was removed by centrifugation (20.000 rpm, 20 min, 4°C), and the pellet was resuspended in 400 μl two-fold concentrated SDS PAGE sample buffer and heat-denatured. 50 μl from the supernatant containing the soluble nucleoplasm was mixed with five-fold concentrated SDS PAGE sample buffer and heat-denatured.

### Growth assays

To analyze the sensitivity of HeLa control and FAM104A/B double knockout cells towards inhibition of p97 and camptothecin, 1000 cells per cell line were seeded in CellCarrier-96 Ultra microplates. 20 h post seeding, half of the wells were treated with 1 uM camptothecin for 70 min. After removal of the inhibitor and two washing steps with PBS, cells in all wells were grown in the absence or presence of CB-5083 (0 – 200 nM) for further seven days. Media containing the indicated CB-5083 concentrations were refreshed every 1.5 days. The experiment was performed in two biological replicates with three technical replicates each. After a total of eight days, cells were washed with PBS, fixed using 3.7% formaldehyde in PBS for 12 min at RT, and washed twice with PBS.

Permeabilization and Hoechst staining was performed by adding 0.2% Triton X-100 in PBS with 2.5 μg/ml Hoechst 33342 for 10 min in the dark. After washing the wells two times with PBS, images were taken with an Operetta® CLS High-Content Imaging System with 20-fold magnification and analyzed using the Harmony® High-Content and Imaging Analysis Software (PerkinElmer). 26 image fields per well were acquired.

### Statistical testing

All statistical analyses and graphing were performed using GraphPad Prism 9. Relative cell viability was calculated by normalizing the cell number of each condition to the respective mean cell number of all replicates in the absence of CB-5083, separately for FAM104A/B DKO and control cells. After verifying the assumption of normal distribution using the D’Agostino-Pearson omnibus K^2^ test and excluding a few extreme outliers (less then 3% of the data set), a two-way factorial analysis of variance (ANOVA) was performed. This analysis was followed up by Bonferroni’s multiple comparisons test which corrected for multiple comparisons using statistical hypothesis testing. A p value below 0.05 was considered significant and indicated as following: ns = not significant, * p < 0.05, ** p < 0.01.

## Acknowledgements

We thank Manuel Kaulich and Didier Trono for providing plasmids; the Imaging Core Facility (Biocenter, University of Würzburg) for support with confocal microscopy; and members of the Buchberger lab for critical reading of the manuscript. This work was funded by the Deutsche Forschungsgemeinschaft (DFG, German Research Foundation) through grants GRK2243/1+2 (to A.B.) and 440766788 (INST 93/1023-1-FUGG; Operetta CLS system).

## Author Contributions

M.Kö., G.M. and A.B. conceived the project; M.Kö., S.M., G.M., M.Ke. and C.S.-V. performed experiments; C.G. and K.H. performed structural and bioinformatic analyses; M.Kö., G.M., M.Ke., U.F. and A.B. analyzed the data; M.Kö. and A.B. wrote the manuscript with support from all other authors.

## Additional information

Correspondence and requests for materials should be addressed to A.B.

## Supplementary Figures

**Figure S1.**
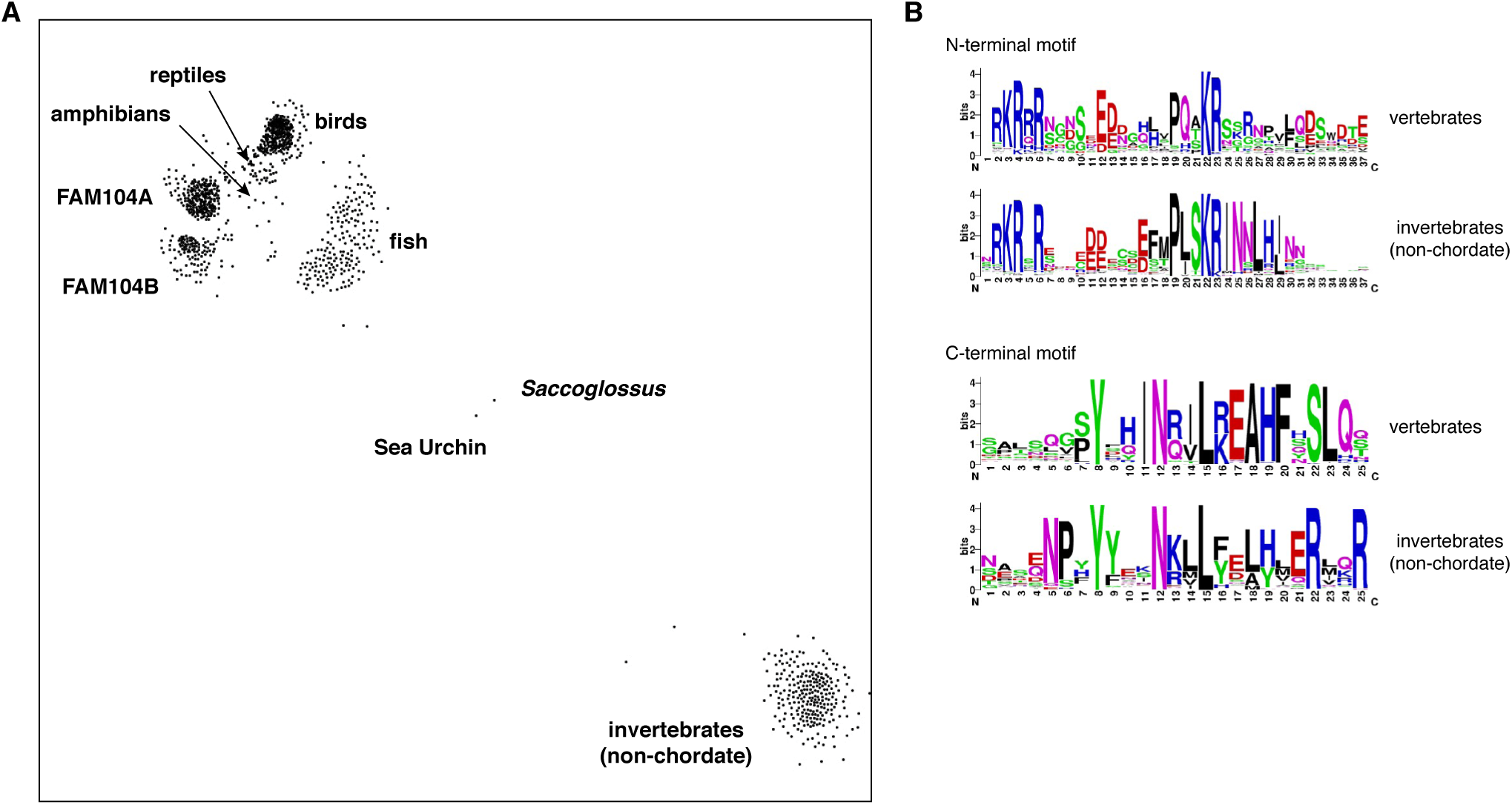
Evolutionary conservation of FAM104 proteins. **(A)** CLANS diagram of the FAM104 family. The sequence relationship between different FAM104-related proteins is visualized by similarity-based clustering, using the CLANS software. Each point represents one protein; point-to point distances are approximations of sequence divergence. Selected clusters are annotated. **(B)** Consensus sequences of conserved motifs. Separate sequence logos for the N- and C-terminal motifs were generated for the vertebrate and invertebrate clusters, respectively.

**Figure S2.**
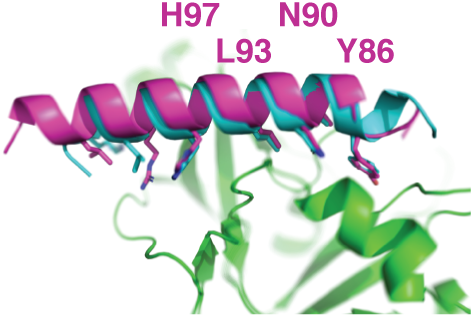
The p97 binding mode is conserved in invertebrate FAM104 homologs. AlphaFold Multimer model of the C-terminal alpha-helix of *Drosophila melanogaster* CG14229 (purple) superimposed on the model of FAM104A (turquoise) bound to the N domain of p97 (green) shown in Fig. 2G. The modeled N domain of *Drosophila melanogaster* TER94 was almost identical to that of p97 and was omitted for clarity. The four highly conserved residues contacting the N domain are labeled.

**Figure S3.**
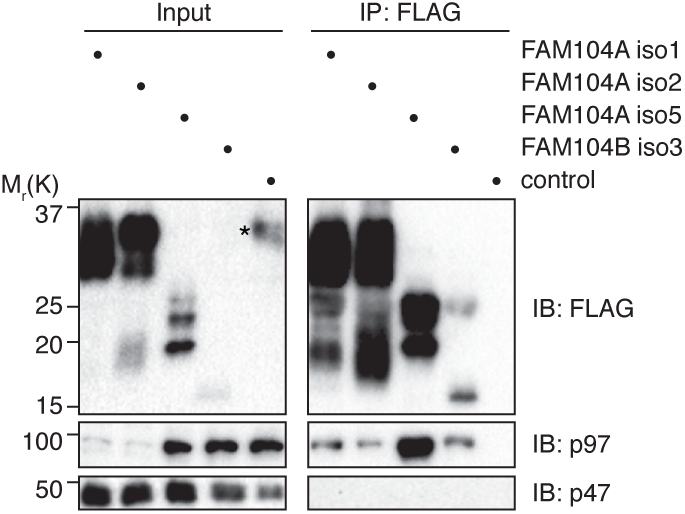
p47 does not co-immunoprecipitate with FAM104 proteins. Immunoprecipitation (IP) experiment as in Fig. 3A. Input and IP samples were immunoblotted for the FLAG epitope tag, p97, and p47. In the input lane of the control sample, an asterisk in the FLAG immunoblot labels a spill-over from the strong signal in the neighboring IP lane.

**Figure S4.**
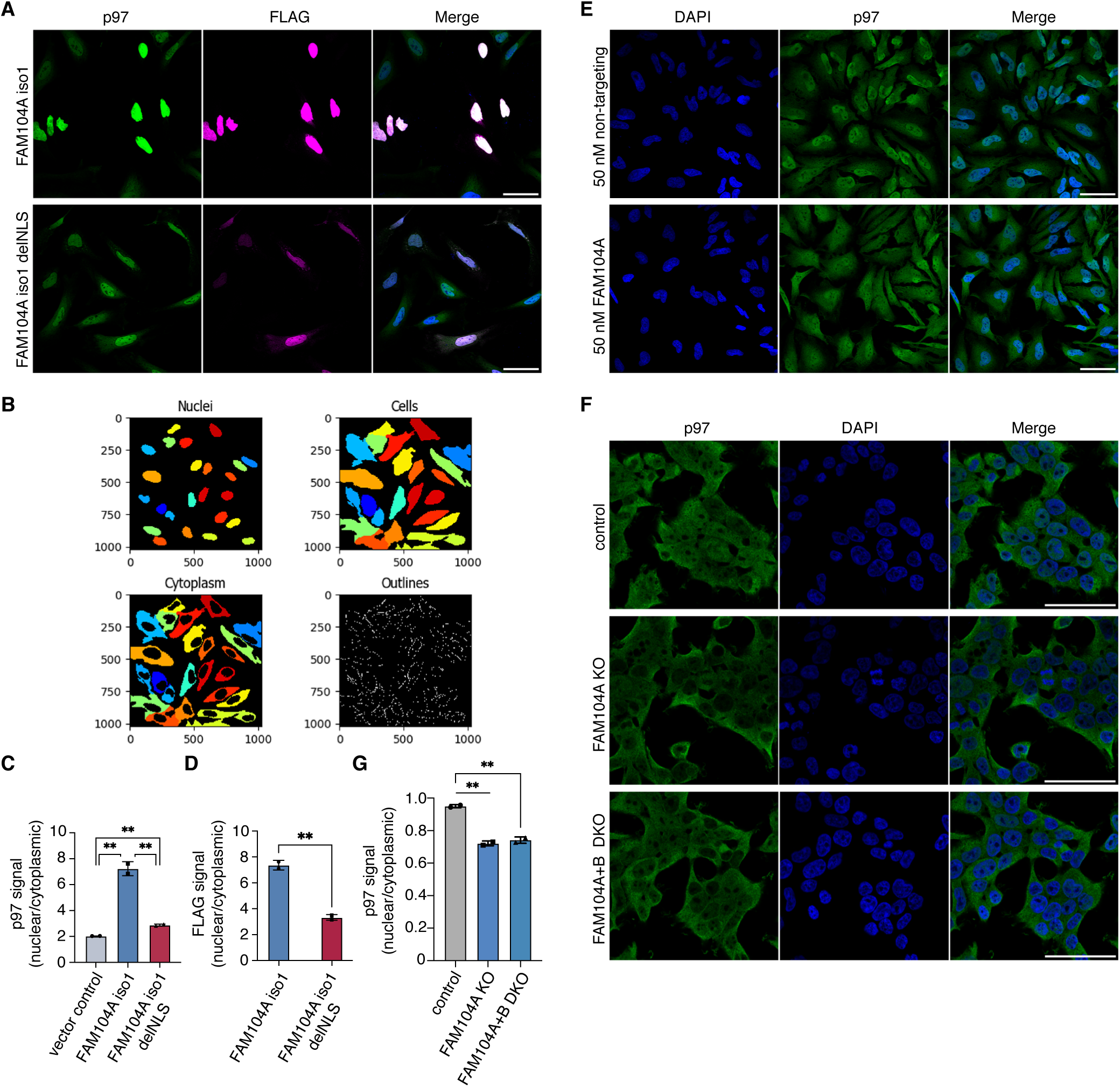
FAM104A/B promote the nuclear localization of p97. **(A)** Comparison of the images from Fig. 4AB for full-length FAM104A isoform 1 and its cNLS-deleted variant. The images were processed for higher signal intensities in order to facilitate the visual inspection of cytoplasmic signals. Scale bars, 50 µm. **(B)** Example of the cell segmentation results for nuclear and cytoplasmic signals obtained with CellProfiler. See Methods section for details. **(C, D)** Quantification of the ratios of nuclear to cytoplasmic p97 (C) and FLAG (D) signals detected in (A). For the analysis of FAM104 isoform 1 and its cNLS-deleted variant, only transfected cells (as judged by the FLAG channel) were included in the quantification. Shown is the mean ± s.d.; n = 2 with 15 -30 transfected cells per replicate and condition; **, p < 0.01; Student’s t-test. **(E)** Z stack images of control- or FAM104A-depleted HeLa cells stained for endogenous p97. Shown are maximum intensity projections of stacks consisting of eight z planes with 2.5 µm distance. **(F)** Control, FAM104A single knockout (KO) and FAM104A/FAM104B double knockout (DKO) HEK293T cell pools were analyzed by confocal immunofluorescence microscopy, using an antibody against endogenous p97. **(G)** Quantification of nuclear to cytoplasmic p97 signals detected in (F). Scale bars, 50 µm

**Supplementary Table S1.**

| Reagent or resource | Source | Identifier |
| --- | --- | --- |
| **Antibodies** |  |  |
| Mouse monoclonal anti-alpha-tubulin | Sigma-Aldrich | Cat# T5168, RRID: AB_477579 |
| Mouse monoclonal anti GAL4-TA | Santa Cruz | Sc-1663 |
| Rabbit polyclonal anti-FLAG | Thermo Fisher Scientific | Cat# PA1-984B, RRID: AB_347227 |
| Mouse monoclonal anti-FLAG | Sigma-Aldrich | Cat# F1804 |
| Rabbit polyclonal anti-VCP | Bethyl Laboratories | Cat# A300-589A, RRID: AB_495512 |
| Mouse monoclonal anti-VCP | Santa Cruz | Cat# sc-57492, RRID: AB_793927 |
| Mouse monoclonal anti-VCP Alexa Fluor™ 647 conjugate | Santa Cruz | Cat# sc-57492 |
| Rabbit polyclonal anti-FAF1 | Max Planck Institute of Biochemistry, animal house | AB65 |
| Rabbit polyclonal anti-UBXN7 | Sigma-Aldrich | Cat# HPA049442 |
| Rabbit polyclonal anti-NPL4 | Sigma-Aldrich | Cat# HPA021560 |
| Rabbit polyclonal anti-UFD1 | Proteintech | Cat# 10615 |
| Rabbit monoclonal anti-Ubiquityl-Histone H2B | Cell Signaling | Cat# 5546 |
| Goat polyclonal anti-Mouse IgG HRP | Dianova (Jackson ImmunoResearch) | Cat# 115-035-003, RRID:AB_10015289 |
| Goat polyclonal anti-Rabbit IgG HRP | Dianova (Jackson ImmunoResearch) | Cat# 111-035-045, RRID: AB_2337938 |
| Alexa Fluor 488 Goat Anti-Rabbit IgG (H+L) | Thermo Fisher Scientific | Cat# A-11070, RRID: AB_142134 |
| Alexa Fluor 488 Goat Anti-Mouse IgG (H+L) | Thermo Fisher Scientific | Cat# A-11017, RRID: AB_143160 |
| Alexa Fluor 594 Goat Anti-Rabbit IgG (H+L) | Thermo Fisher Scientific | Cat# A-11072, RRID: AB_142057 |
| Alexa Fluor 594 Goat Anti-Mouse IgG (H+L) | Thermo Fisher Scientific | Cat# A-11020, RRID: AB_141974 |
| **Bacterial strains** |  |  |
| XL1 Blue | Stratagene | Cat# 200249 |
| XL10 Gold | Agilent | Cat# 200516-4 |
| BL21(DE3) pRIL | Agilent | Cat# 230245 |
| **Yeast strain** |  |  |
| PJ69-4A | (James *et al*, 1996) | N/A |
| **Critical commercial resources** |  |  |
| Camptothecin | Selleckchem | NSC-100880 |
| CB-5083 | Selleckchem | Cat# S8101 |
| Clarity Western ECL Substrate | Bio-Rad | Cat# 1705061 |
| cOmplete, EDTA-free Protease Inhibitor Cocktail | Roche | Cat# 04693132001 |
| Glutathione Sepharose 4 Fast Flow | Cytiva | Cat# 17513202 |
| Ni-NTA Agarose | Qiagen | Cat# 30230 |
| Opti-MEM | Thermo Fisher Scientific | Cat# 31985602 |
| Polybrene | Santa Cruz | Cat# sc-134220 |
| Polyethylenimine (PEI) | Polysciences | Cat# 23966-1 |
| ProLong Glass Antifade Mountant | Thermo Fisher Scientific | Cat# P36980 |
| Puromycin | InvivoGen | Cat# ant-pr-1 |
| Anti FLAG-M2 affinity agarose beads | Sigma-Aldrich | Cat# A2220 |
| Oligofectamine™ | Thermo Fisher Scientific | Cat# 12252011 |
| Pierce™ BCA Protein Assay Kit | Thermo Fisher Scientific | Cat# 23225 |
| QuickChange XLII Mutagenesis Kit | Agilent | Cat# 200521 |
| **Experimental models: cell lines** |  |  |
| HeLa | ATCC | CCL-2 |
| HEK293T | ATCC | CRL-3216 |
| HeLa FAM104A knockout cell pool | This study | N/A |
| HeLa FAM104A+B double knockout cell pool | This study | N/A |
| HeLa non-human-target control cell pool | This study | N/A |
| HEK293T FAM104A knockout cell pool | This study | N/A |
| HEK293T FAM104A+B double knockout cell pool | This study | N/A |
| HEK293T non-human-target control cell pool | This study | N/A |
| **Oligonucleotides** |  |  |
| ON-TARGETplus human FAM104A siRNA - SMARTpool | Dharmacon | Cat# L-015015-02-0005 |
| ON-TARGETplus Non-targeting pool | Dharmacon | Cat# D-001810-10-05 |
| **Recombinant DNA** |  |  |
| psPAX2 | Addgene | Cat# 12260 |
| pMD2.G | Addgene | Cat# 12259 |
| pCMV-Tag2B | Agilent | Cat# 211172 |
| pCMV FAM104A isoform 1 | This study | pAB2225 |
| pCMV FAM104A isoform 2 | This study | pAB2208 |
| pCMV FAM104A isoform 5 | This study | pAB2223 |
| pCMV FAM104A isoform 1 Cdel26 | This study | pAB2280 |
| pCMV FAM104A isoform 2 Cdel26 | This study | pAB2275 |
| pCMV FAM104A isoform 5 Cdel26 | This study | pAB2276 |
| pCMV FAM104A isoform 1 delNLS | This study | pAB2230 |
| pCMV FAM104A isoform 2 delNLS | This study | pAB2231 |
| pCMV FAM104A isoform 5 delNLS | This study | pAB2227 |
| pGEX-4T1 | Cytiva | Cat# 28954549 |
| pGEX-4T1 FAM104A isoform 1 | This study | pAB2212 |
| pGEX-4T1 FAM104A isoform 2 | This study | pAB2195 |
| pGEX-4T1 FAM104A isoform 5 | This study | pAB2213 |
| pGEX-4T1 FAM104B isoform 3 | This study | pAB2298 |
| pGEX-4T1 FAM104A isoform 1 Cdel26 | This study | pAB2295 |
| pGEX-4T1 FAM104A isoform 2 Cdel26 | This study | pAB2296 |
| pGEX-4T1 FAM104A isoform 5 Cdel26 | This study | pAB2297 |
| pGEX-4T1 FAM104B isoform 3 Cdel26 | This study | pAB2299 |
| pGEX-4T1 FAM104A isoform 5 NL->AA | This study | pAB3163 |
| pGEX-4T1 FAM104A isoform 5 NL->RR | This study | pAB3162 |
| mini-pRSETA | (Perrett *et al*, 1999) | N/A |
| mini-pRSETA UBXN7 | This study | pAB2008 |
| mini-pRSETA p47 | (Allen *et al*, 2006) | pAB356 |
| pQE30 FAF1 | (Jensen *et al*, 2001) | N/A |
| pET28a(+) UFD1 | (Fernandez-Saiz & Buchberger, 2010) | pAB425 |
| pET21d NPL4 | (Fernandez-Saiz & Buchberger, 2010) | pAB1340 |
| pProExHT p97 | (Fernandez-Saiz & Buchberger, 2010) | pAB1312 |
| pProExHT p97 N domain | (Fernandez-Saiz & Buchberger, 2010) | pAB1342 |
| pProExHT p97 ND1 | (Fernandez-Saiz & Buchberger, 2010) | pAB1343 |
| pProExHT p97 ΔN | (Rothballer *et al*, 2007) | pAB749 |
| pGAD-C1 | (James *et al*, 1996) | N/A |
| pGAD-C1 FAM104A isoform 1 | This study | pAB2217 |
| pGAD-C1 FAM104A isoform 2 | This study | pAB2205 |
| pGAD-C1 FAM104A isoform 5 | This study | pAB2215 |
| pGAD-C1 FAM104B isoform 3 | This study | pAB2233 |
| pGBDU | (James *et al*, 1996) | N/A |
| pGBDU p97 | This study | pAB1184 |
| pGAD-C1 FAM104A isoform 5 Cdel4 | This study | pAB2312 |
| pGAD-C1 FAM104A isoform 5 Cdel7 | This study | pAB2313 |
| pGAD-C1 FAM104A isoform 5 Cdel13 | This study | pAB2314 |
| pGAD-C1 FAM104A isoform 5 Cdel18 | This study | pAB2315 |
| pGAD-C1 FAM104A isoform 5 Cdel26 | This study | pAB2235 |
| pLentiCRISPRv2 | Addgene | Cat# 52961 |
| Non human control sgRNAs in pLentiCRISPRv2 | Manuel Kaulich, University of Frankfurt | N/A |
| FAM104A sgRNAs in pLentiCRISPRv2 | Manuel Kaulich, University of Frankfurt | N/A |
| FAM104B sgRNAs in pLentiCRISPRv2 | Manuel Kaulich, University of Frankfurt | N/A |
| **Software** |  |  |
| Fiji | http://fiji.sc | RRID: SCR_002285 |
| Image Lab Software | Bio-Rad | RRID: SCR_014210 |
| CellProfiler | https://cellprofiler.org/ |  |
| GraphPad Prism | https://www.graphpad.com/scientific-software/prism/ |  |

